## Supplementary material for "Modification-Dependent Restriction Endonuclease-based sequencing method (EcoWI-seq) maps the genome-wide landscape of phosphorothioate modification at base resolution": High quality Figures and Supplementary Figures

### Figure 1

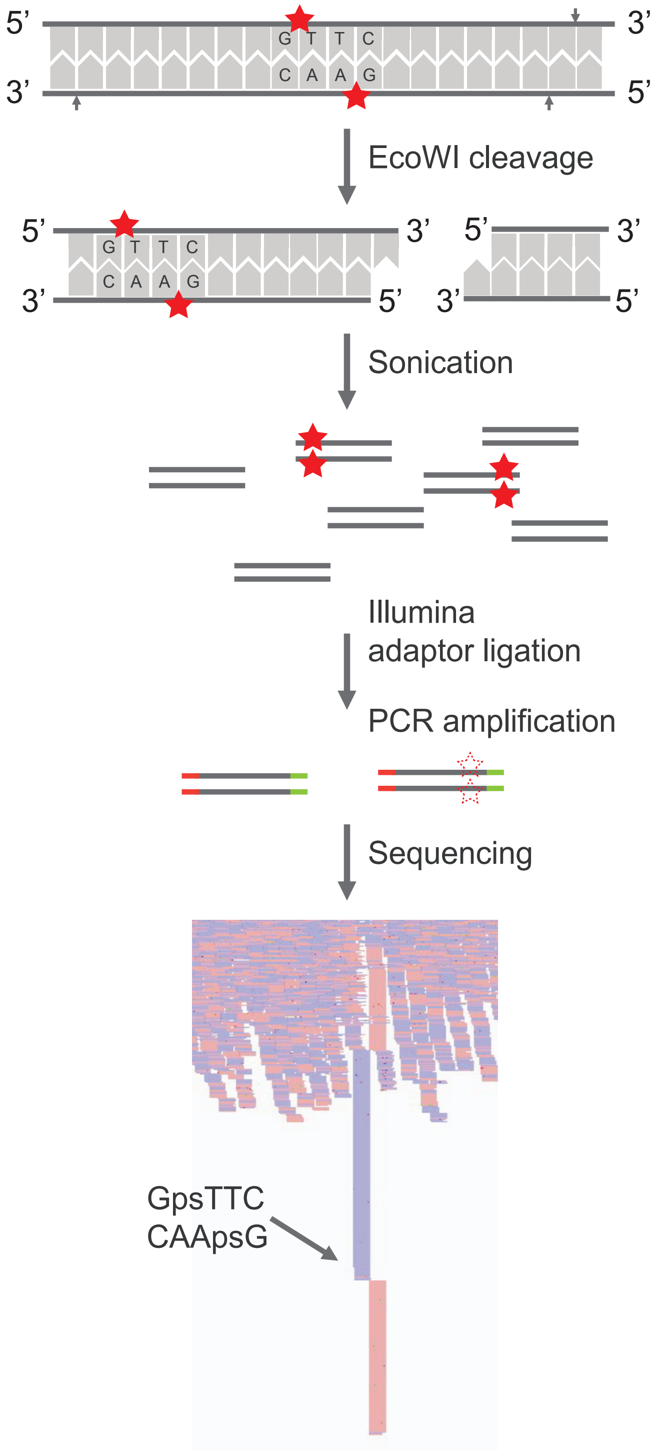
5’
3’
G T T C
C A A G
3’
5’
EcoWI cleavage
5’
3’
5’
3’
G T T C
C A A G
3’
5’
3’
5’
Sonication
Illumina
adaptor ligation
PCR amplification
Sequencing
GpsTTC
CAApsG

### Figure 2

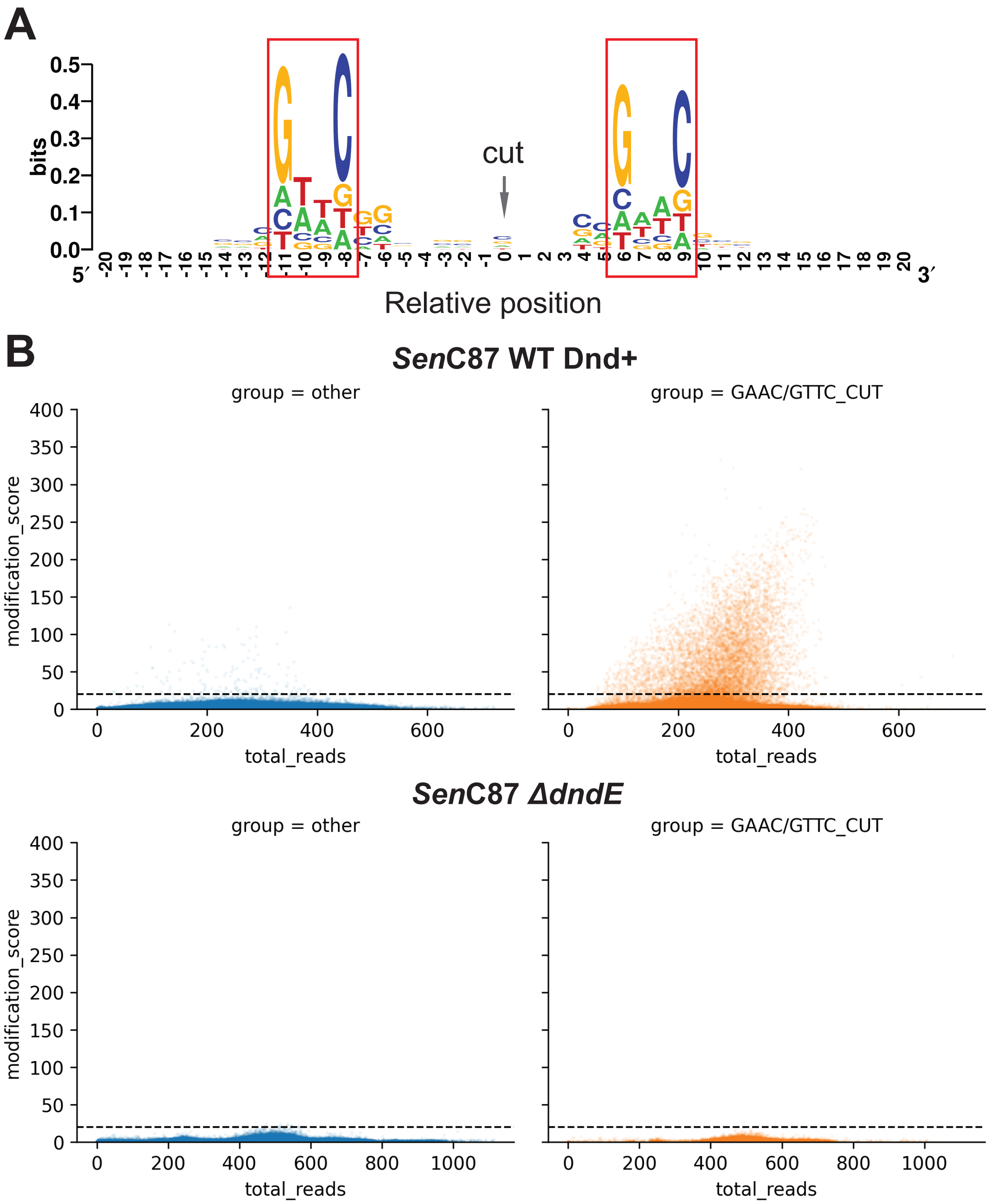


C

| | Total sites | GAAC/GTTC | Other |
|---|---|---|---|
| ***Sen*C87 WT** | 3973 | 3870 | 103 |
| ***Sen*C87 *ΔdndE*** | 1 | 0 | 1 |

Figure 3

**A**

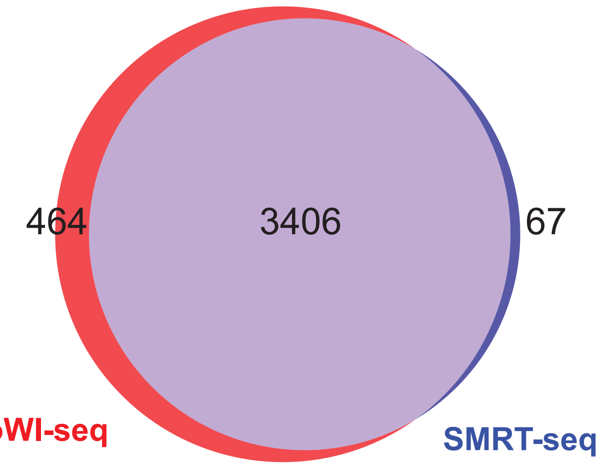


| EcoWI-seq sites | SMRT-seq sites | Overlap sites |
|---|---|---|
| 3870 | 3473 | 3406 (98.1%) |
| 3870 | 3473 (random) | 424 (12.2%) |

**B**

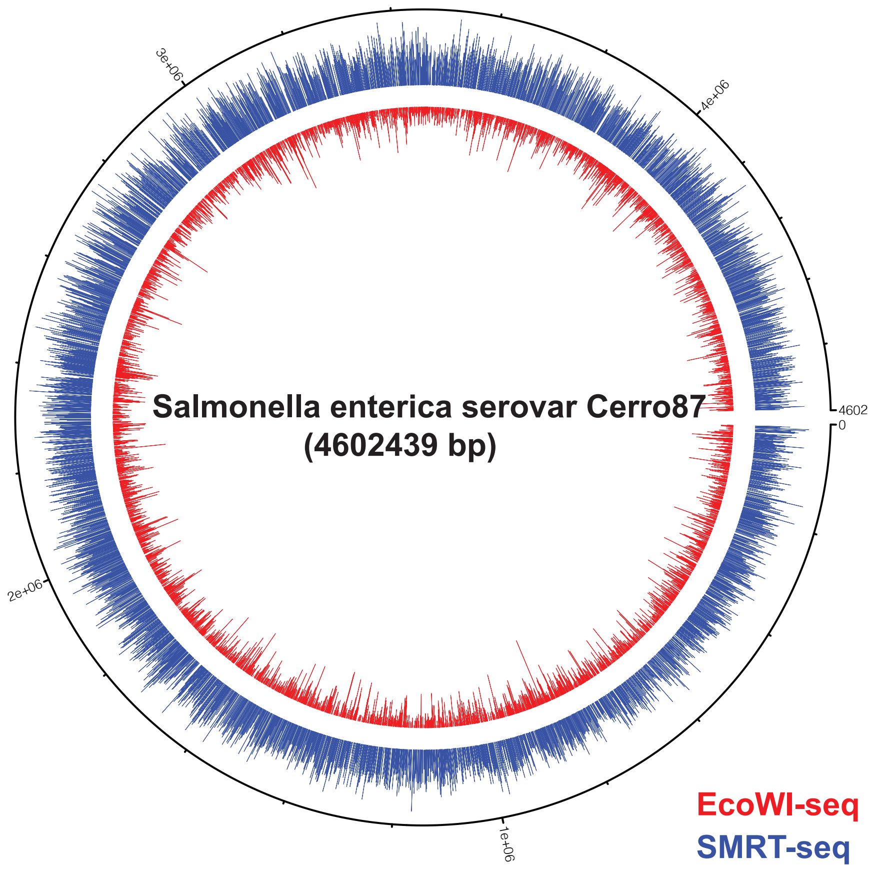

Figure 4

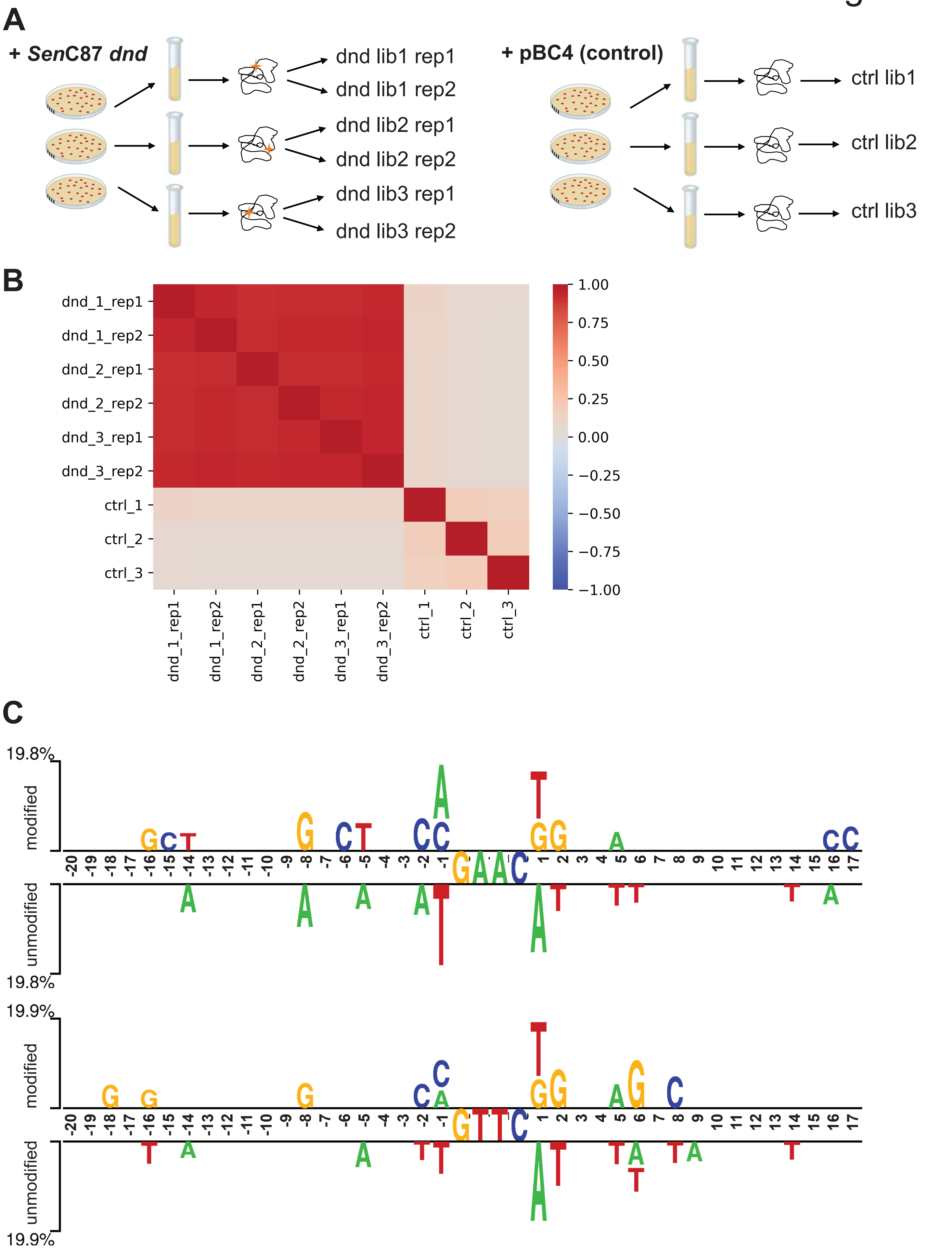
A
+ SenC87 dnd
dnd lib1 rep1
dnd lib1 rep2
dnd lib2 rep1
dnd lib2 rep2
dnd lib3 rep1
dnd lib3 rep2
+ pBC4 (control)
ctrl lib1
ctrl lib2
ctrl lib3
B
dnd_1_rep1
dnd_1_rep2
dnd_2_rep1
dnd_2_rep2
dnd_3_rep1
dnd_3_rep2
ctrl_1
ctrl_2
ctrl_3
1.00
0.75
0.50
0.25
0.00
−0.25
−0.50
−0.75
−1.00
C
19.8%
modified
unmodified
19.8%
19.9%
modified
unmodified
19.9%

### Supplementary Figure S1

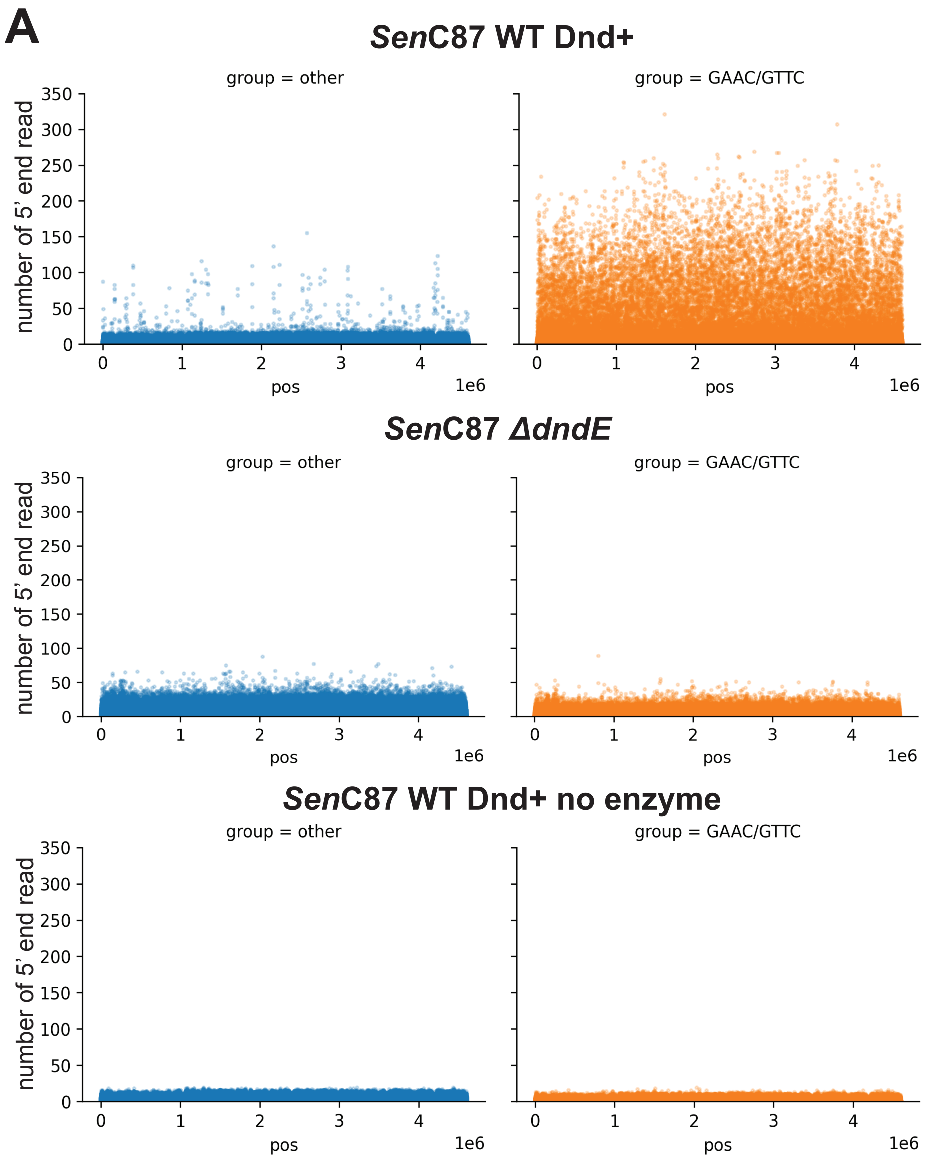


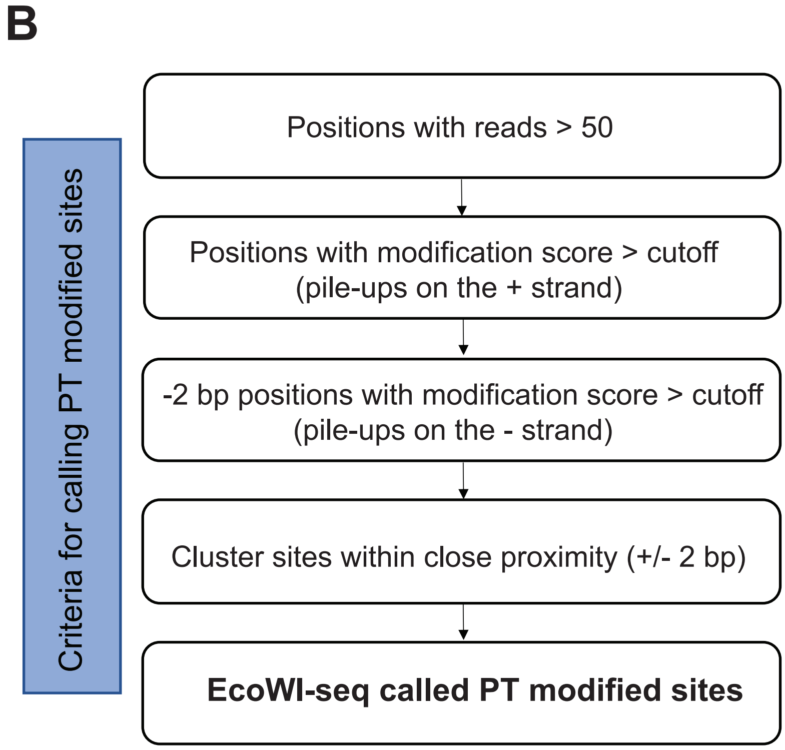

### Supplementary Figure S2

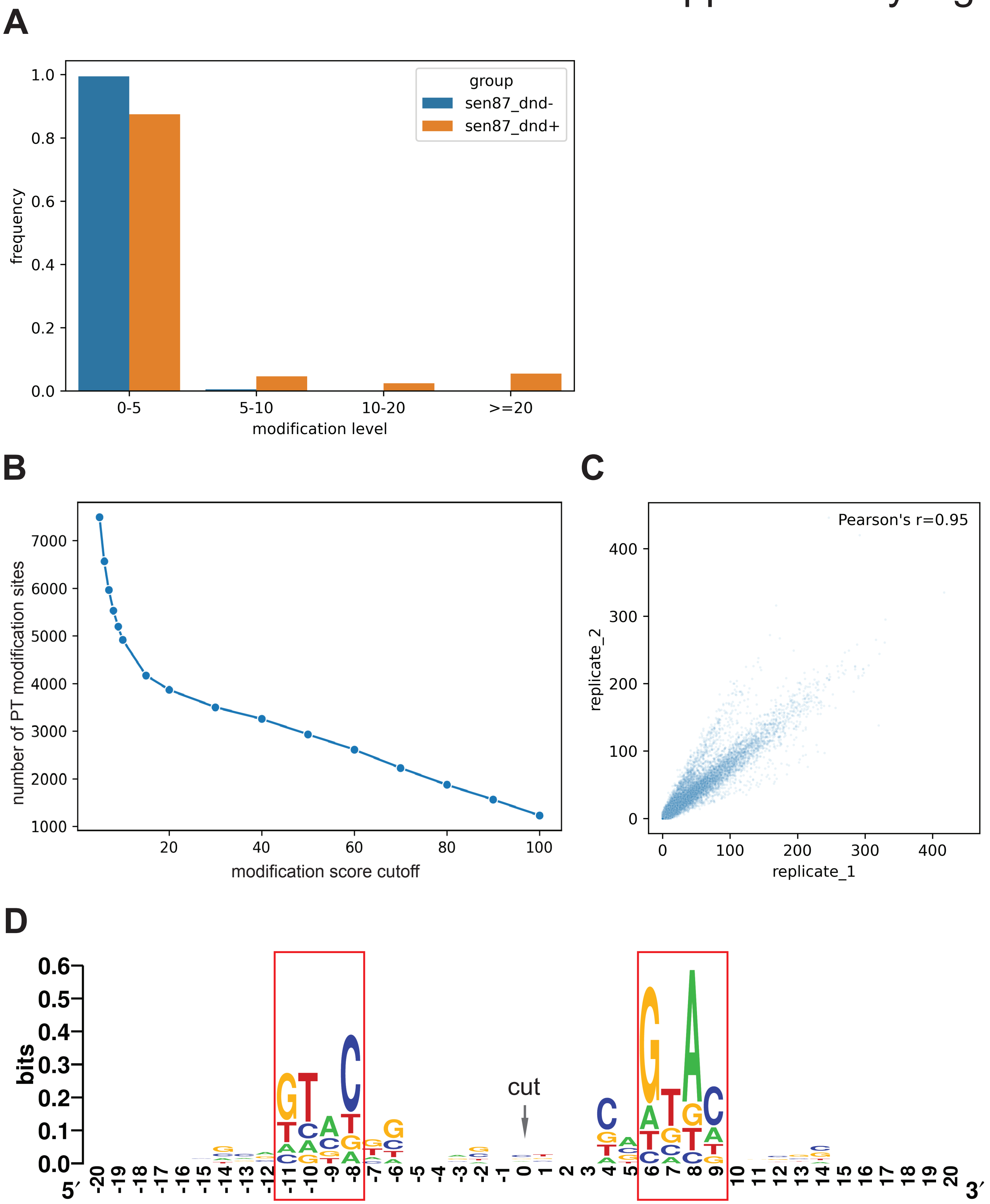

### Supplementary Figure S3

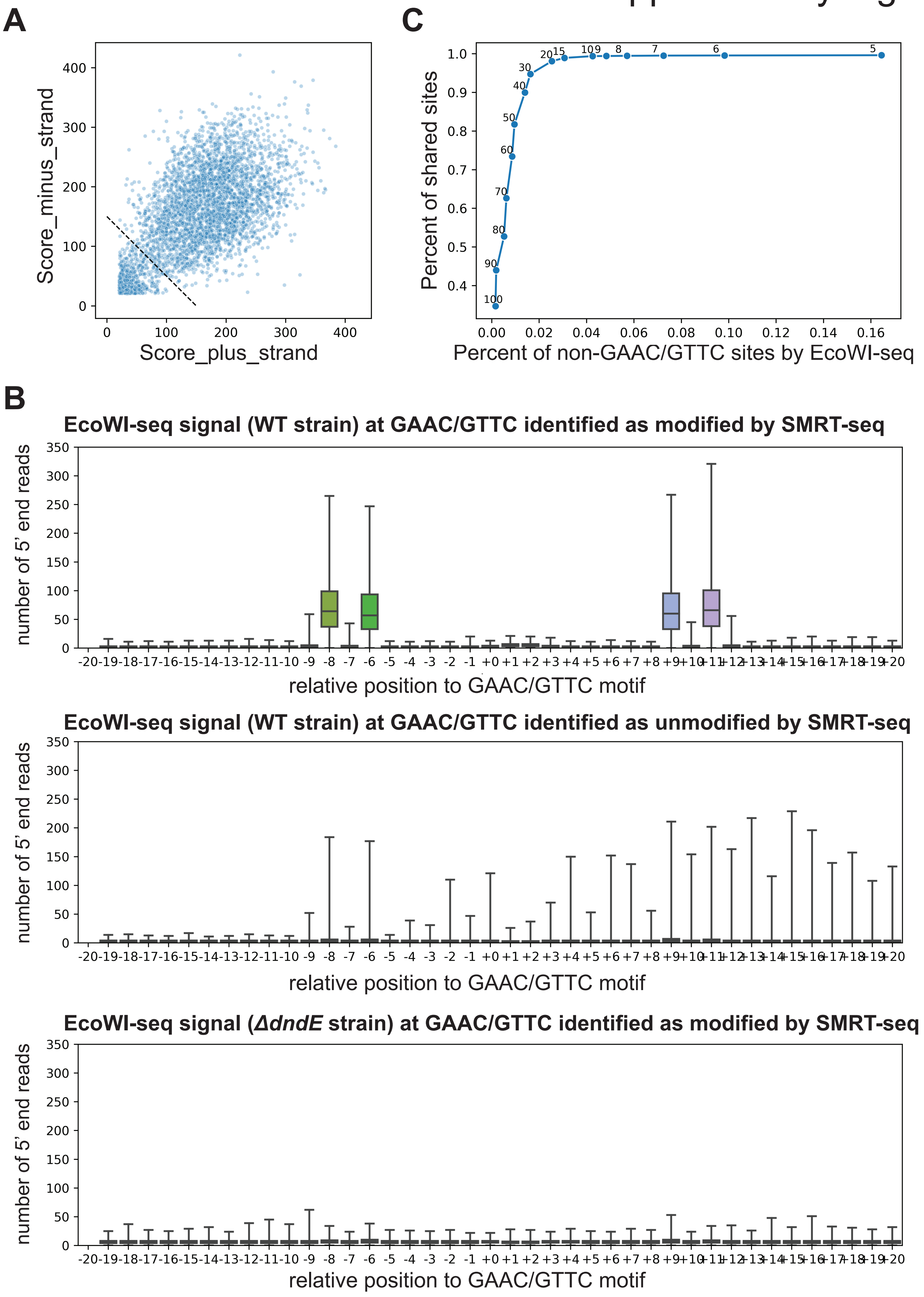

### Supplementary Figure S4

**A**

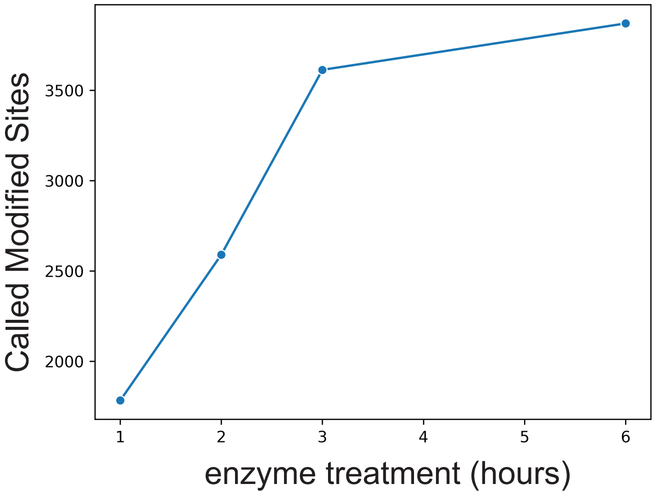


**B**

| Incubation | EcoWI-seq sites | Overlap sites with SMRT-seq | SMRT percent | EcoWI percent |
|---|---|---|---|---|
| 6 hours | 3870 | 3407 | 98.1% | 88.0% |
| 3 hours | 3613 | 3217 | 92.6% | 89.0% |
| 2 hours | 2590 | 2527 | 72.8% | 97.6% |
| 1 hours | 1783 | 1729 | 49.8% | 97.0% |

**C**

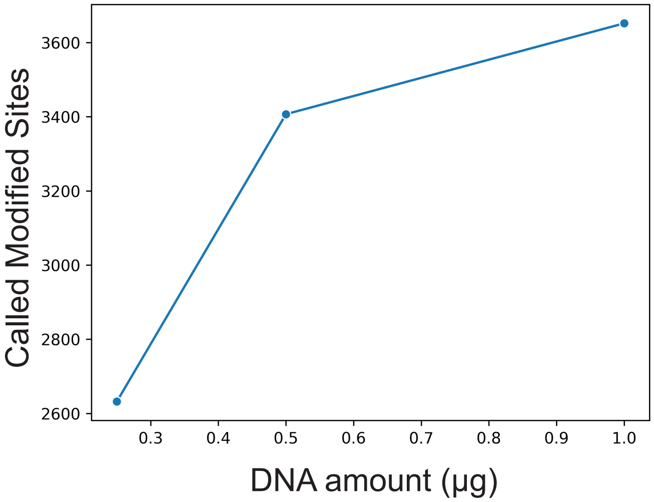


**D**

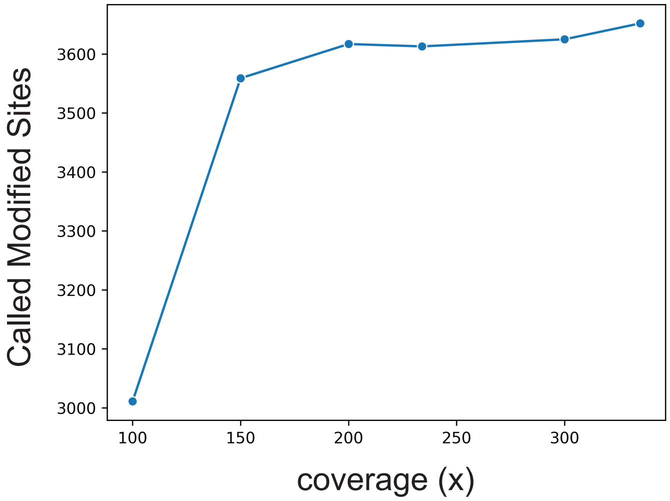


**E**

| Sequencing | EcoWI-seq PT sites | Overlap sites with SMRT-seq | Overlap percent |
|---|---|---|---|
| single read | 3642 | 3221 | 92.7% |
| paired reads | 3609 | 3203 | 92.3% |

Supplementary Figure S5

**A**

| | Assembled genome |
|---|---|
| span (bp) | 4,548,135 |
| N (%) | 0.00 |
| GC (%) | 52.29 |
| AT (%) | 47.71 |
| scaffold count | 252 |
| longest scaffold (bp) | 294,596 |
| scaffold N50 length (bp) | 95,113 |
| scaffold N50 count | 15 |
| scaffold N90 length (bp) | 27,362 |
| scaffold N90 count | 47 |
| contig count | 252 |
| contig N50 length (bp) | 95,113 |
| contig N50 count | 15 |
| contig N90 length (bp) | 27,362 |
| contig N90 count | 47 |

**B**

| Reference genome | De novo assembly | Overlap sites |
|---|---|---|
| 3973 | 3833 | 3741 (97.6%) |

**A**

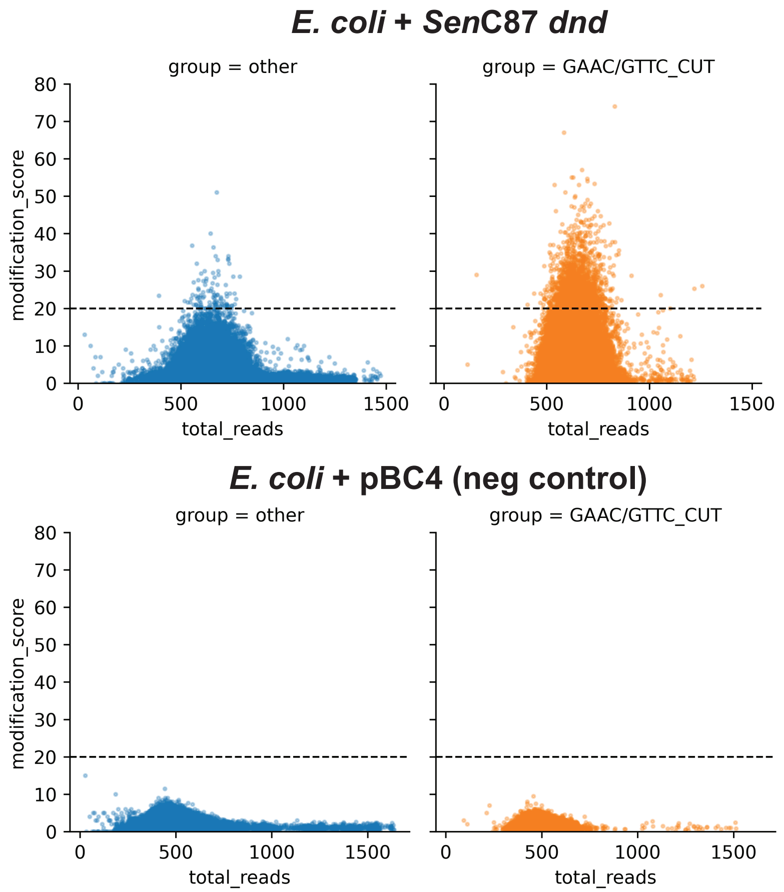


**B**

| | Total sites | GAAC/GTTC | Other |
|---|---|---|---|
| ***E.coli* + *SenC87 dnd*** | 1696 | 1616 | 80 |
| ***E.coli* + pBC4** | 0 | 0 | 0 |

**C**

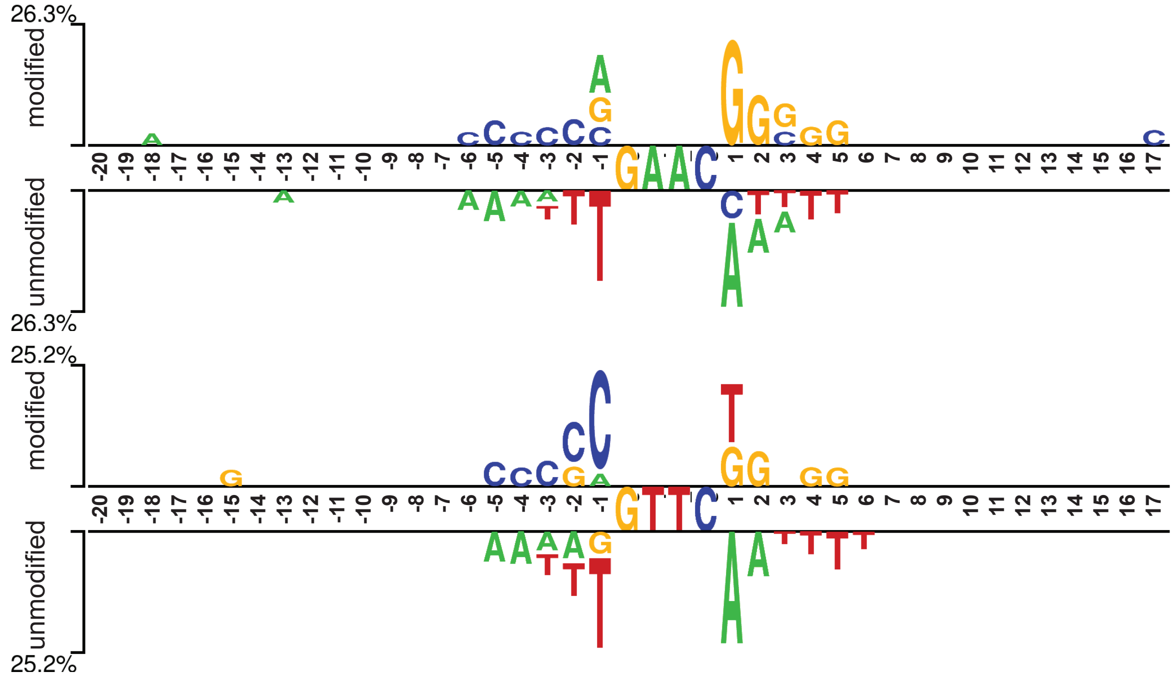
